# Prime editing enables precise genome editing in mouse liver and retina

**DOI:** 10.1101/2021.01.08.425835

**Authors:** Hyewon Jang, Jeong Hong Shin, Dong Hyun Jo, Jung Hwa Seo, Goosang Yu, Ramu Gopalappa, Sung-Rae Cho, Jeong Hun Kim, Hyongbum Henry Kim

## Abstract

Prime editing can induce any small-sized genetic change without donor DNA or double strand breaks. However, it has not been investigated whether prime editing is possible in postnatal animals. Here we delivered prime editors 2 and 3 into a mouse model of hereditary tyrosinemia, a genetic liver disease, using hydrodynamic injection, which corrected the disease-causing mutation and rescued the phenotype. We also achieved prime editing in the retina and retina pigment epithelium in wild-type mice by delivering prime editor 3 using trans-splicing adeno-associated virus. Deep sequencing showed that unintended edits at or near the target site or off-target effects were not detectable except for low levels (0% to 1.2%) of indels when PE3, but not PE2, was used. Our study suggests that precise, prime editor-mediated genome editing is possible in somatic cells of adult animals.

---

Prime editing can induce any small-sized genetic change, including insertions, deletions, and all 12 possible point mutations, without donor DNA or double strand breaks^1^. There are four types of prime editors (PEs): PE1, PE2, PE3, and PE3b^1^. PE1 and PE2 are composed of a Cas9 nickase-reverse transcriptase (RT) fusion protein and a prime editing guide RNA (pegRNA)^1^. PE3 and PE3b consist of PE2 and one additional single guide RNA (sgRNA)^1^.

Prime editing has been used in cultured mammalian cells^1, 2^ and organoids^3^, plants^4^, and mouse embryos^5^ to introduce genetic changes in a targeted manner. However, the efficiency and precision of prime editing in postnatal mammals in vivo have not been determined. In this study, we show that prime editing can be induced in the liver and eyes of adult mice and that it can correct a genetic disease in a highly precise manner.

## Results

### A mouse model of tyrosinemia and high-throughput evaluation of pegRNAs

As a prototypic disease model for prime editing-based therapeutic genome editing in adult animals, we chose a mouse model of hereditary tyrosinemia type 1 (*Fah*^*mut/mut*^), which is caused by a loss-of-function mutation in the fumarylacetoacetate hydrolase (*FAH*) gene^6,7^. The mouse model contains a homozygous G-to-A point mutation at the last nucleotide of exon 8, which causes exon 8 skipping and results in loss-of-function of FAH (Fig. 1a). Pharmacological inhibition of 4-hydroxyphenylpyruvate dioxygenase, an enzyme that acts upstream of FAH in the tyrosine catabolic pathway, with 2-(2-nitro-4-trifluoromethylbenzoyl)-1,3-cyclohexanedione (NTBC) reduces the accumulation of toxic metabolites and thus prevents hepatic injury^6^. We and others have used this mouse model for testing various in vivo genome editing approaches, including a method based on homology directed repair (HDR)^8, 9^, microhomology-mediated end joining^10^, and base editing^11^. In addition, because hydrodynamic injections have commonly been used to deliver genome editing components for Cas9-directed HDR^8^, microhomology-mediated end joining^10^, and base editing^11^ in this mouse model, use of the same delivery method for prime editors should facilitate a comparison of prime editing with the other genome editing approaches.

**Figure 1.**
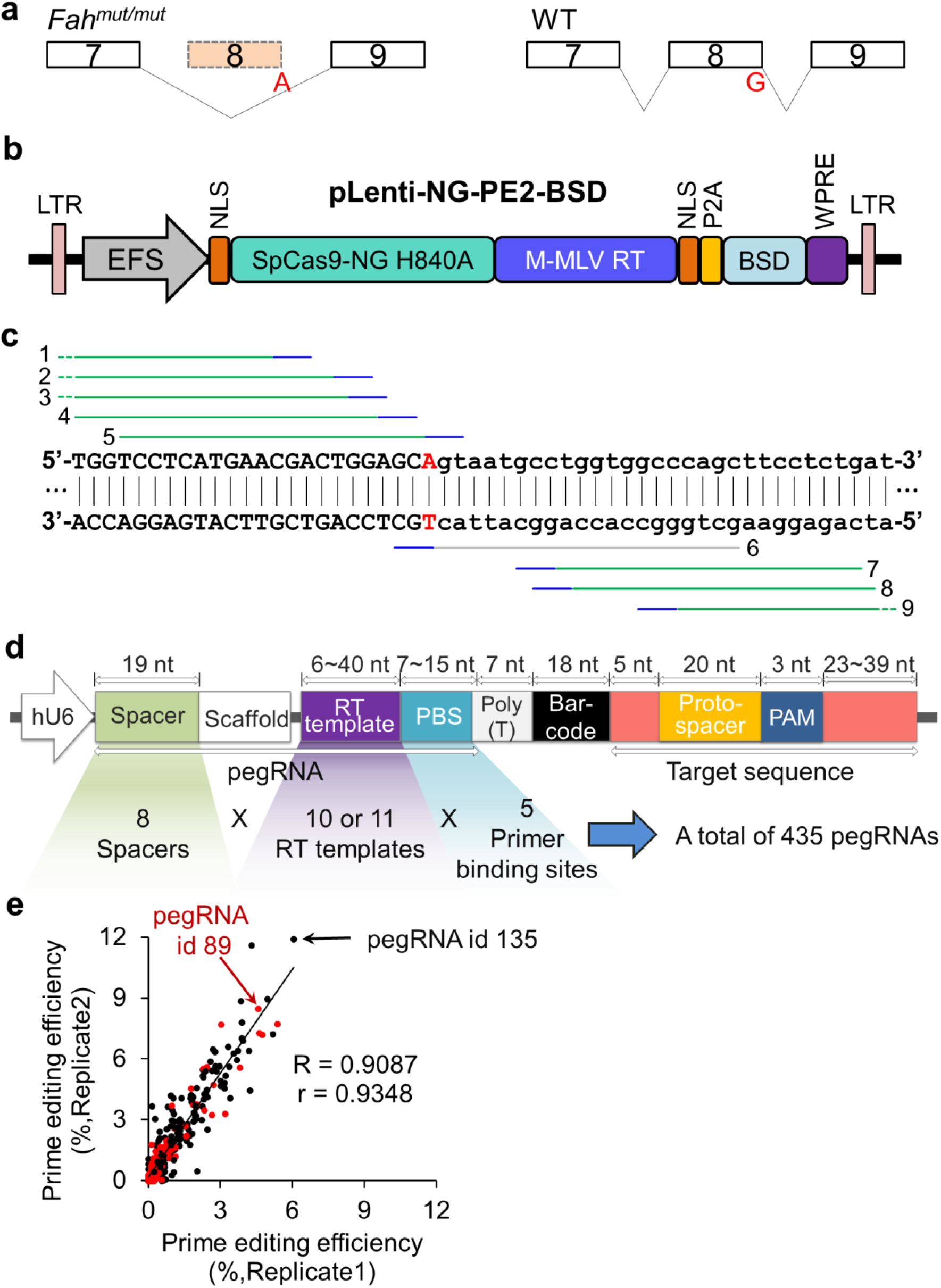
A mouse model of tyrosinemia and high-throughput evaluation of pegRNAs. (**a**) Exon 8 skipping in *Fah*^*mut/mut*^ mice. The G-to-A point mutation (red) at the last nucleotide of exon 8 of the *Fah* gene leads to exon 8 skipping during splicing. (**b**) A schematic representation of the plasmid encoding NG-PE2. NG-PE2, a fusion of SpCas9-NG H840A nickase with Moloney murine leukemia virus reverse transcriptase (M-MLV RT), is expressed from the EFS promoter. The protein encoded by the blasticidin-resistance gene (*BSD*) is co-expressed as a fusion with PE2, from which it is cleaved by the self-cleaving P2A. LTR, long terminal repeat; EFS, elongation factor 1α short promoter; NLS, nuclear localization sequence; P2A, porcine teschovirus-1 2A; WPRE, woodchuck hepatitis virus posttranscriptional regulatory element. (**c**) SpCas9-NG-based PE2 target sequences for correction of the disease-causing point mutation in *Fah*. The protospacer and PAM of each target sequence are represented with green and blue lines, respectively. The numbers on the left and right indicate the target sequence ids. Exon and intron sequences are shown in upper and lower case letters, respectively. A schematic representation of the lentiviral library of pegRNA-target sequence pairs. Each pegRNA is paired with a wide target sequence that includes a protospacer, a PAM, and neighboring sequences. pegRNA expression is driven by the human U6 promoter (hU6). The library included a total of 435 pegRNA-target pairs, with the pegRNAs containing primer binding sites of five different lengths (7, 9, 11, 13, or 15 nt) and 10 to 11 different RT template lengths (6 to 40 nt). Spacer, guide sequence of pegRNA; RT, reverse transcriptase; PBS, primer binding site. (**e**) Correlation between prime editing efficiencies in replicates independently transfected with the NG-PE2-encoding plasmid. The red and black dots indicate pegRNAs with corresponding target sequences with NGG and NGH PAMs, respectively. The red arrow indicates the pegRNA selected for subsequent experiments and the black arrow indicates the pegRNA that showed the highest activity with NG-PE2. The Spearman (R) and Pearson (r) correlation coefficients are shown. The number of pegRNA and target sequence pairs n = 339.

To find a pegRNA that could induce efficient prime editing to correct the pathogenic point mutation, we first identified PE2 target sequences near the mutation site, which were located at positions ranging from −38 base pairs (bp) to +41 bp from the mutation site. Because only two target sequences with an NGG protospacer adjacent motif (PAM) were available near the mutation site, we generated PE2 based on SpCas9-NG (Fig. 1b), a SpCas9 variant that has wider PAM compatibility than SpCas9^12-14^. We identified nine target sequences for SpCas9-NG-based PE2 (for brevity, hereafter, NG-PE2) for the intended editing of the point mutation (Fig. 1c). Given that Cas9 nuclease activity is one of the most important factors determining PE2 efficiency^2^, we first calculated the predicted nuclease activity of SpCas9-NG at the nine sites using DeepSpCas9-NG^14^. One (target id 6) of the nine target sequences showed a very low predicted activity (Supplementary Table 1), so we removed that target sequence from subsequent pegRNA evaluations. For the remaining eight target sequences, we designed pegRNAs containing primer binding sites of five different lengths [7, 9, 11, 13, or 15 nucleotides (nt)] and 10 to 11 different RT template lengths (6 to 40 nt) (Fig. 1d). To undertake a high-throughput evaluation of the resulting 435 pegRNAs, we generated a lentiviral library containing the 435 pegRNA-encoding sequences paired with the corresponding target sequences^2^ (Supplementary Fig. 1). Next, this lentiviral library was transduced into 293T cells to make cell libraries. These cell libraries were then transiently transfected with plasmids encoding NG-PE2. Five days after the transfection, the target sequences were analyzed by deep sequencing. The highest prime editing efficiencies were observed with a pegRNA (pegRNA id 135) that targets a sequence with an NGT PAM (Fig. 1e, Supplementary Table 2). The second and third highest prime editing efficiencies were also observed with pegRNAs (pegRNA ids 136 and 137) that target the same sequence targeted by pegRNA id 135. The fourth and fifth highest prime editing efficiencies were 6.56% and 6.52%, which were observed with two pegRNAs (pegRNA ids 88 and 89 in Supplementary Table 2) targeting a common sequence (target sequence id 2) with an NGG PAM. However, according to DeepSpCas9-NG^14^, the predicted SpCas9-NG nuclease activity at the NGT PAM-containing target sequence that corresponds to pegRNA id 135 was only 17.0, which was substantially lower than 52.4, the SpCas9 nuclease activity predicted by DeepSpCas9^15^ at the NGG PAM-containing target sequence (Supplementary Table 2). Thus, we expected that the prime editing efficiencies of PE2 with pegRNA id 88 or 89 would be higher than that of NG-PE2 with pegRNA id 135.

### Evaluation of prime editor 2 efficiencies using target sequence-containing cells

We next tested the three pegRNAs that showed the highest editing efficiencies with NG-PE2 (ids 135, 136, and 137) and the three pegRNAs associated with an NGG PAM that showed the highest efficiency with NG-PE2 (ids 88, 89, and 90). We generated HEK293T cells containing the target sequence by transduction of target sequence-containing lentiviral vector (Figure 2a). The cells were transiently transfected with plasmids encoding NG-PE2 and pegRNA 135, 136, or 137 or plasmids encoding PE2 and pegRNA 88, 89, or 90. Deep sequencing showed that the average PE2-directed prime editing efficiencies of pegRNAs 88, 89, and 90 were 17.6%, 18.7%, and 12.9%, respectively, which were higher than the NG-PE2-induced prime editing efficiencies of pegRNAs 135, 136, and 137 (3.9%, 3.6%, and 6.3%, respectively) (Figure 2b). Thus, we chose pegRNA 89, which showed the highest editing efficiency, for subsequent studies.

**Figure 2.**
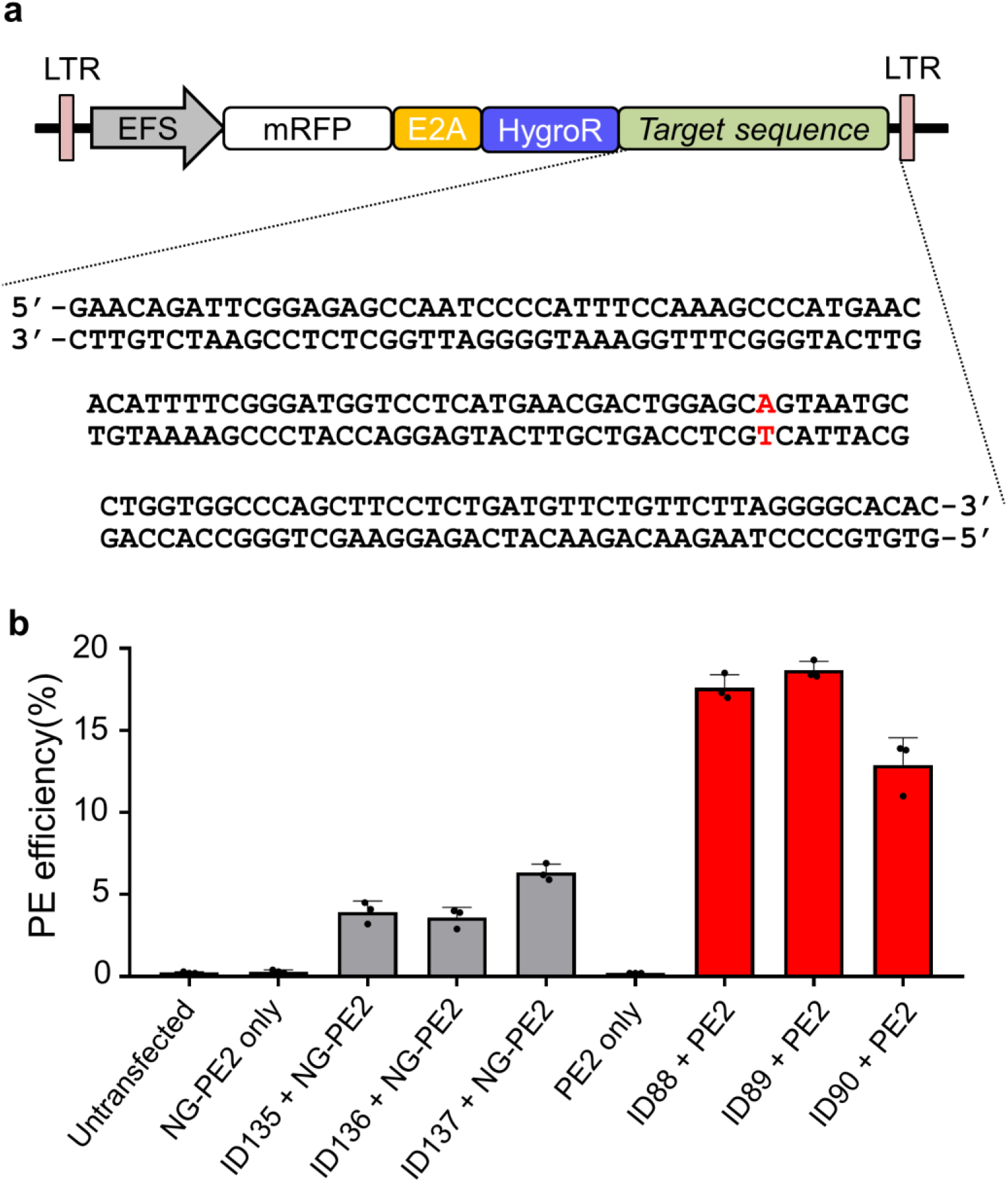
Evaluation of prime editor 2 efficiencies using target sequence-containing cells. (**a**) Schematic representation of the lentiviral vector containing the mutant *Fah* target sequence found in *Fah*^*mut/mut*^ mice. The target sequence is shown; red indicates the mutant base pair. HEK293T cells were transduced with this lentiviral vector to generate target sequence-containing cells. (**b**) Prime editing efficiencies of six pegRNAs (ids 88, 89, 90, 135, 136, and 137) in the target sequence-containing cells. Frequencies were measured five days after transfection of a pegRNA-encoding plasmid and a PE2- or NG-PE2-encoding plasmid. The id of the pegRNA used is shown on the x-axis. Data are mean ± s.d. The number of independent transfections n = 3.

### Prime editor 3 corrects the disease mutation and phenotype in *Fah*^*mut/mut*^ mice

The initial study of prime editing suggested that the editing efficiency of PE3 and PE3b would be, albeit not always, higher than that of PE2 when tested in cultured mammalian cells^1^. Thus, we attempted to use PE3 or PE3b for in vivo genome editing by adding an sgRNA. Based on the DeepSpCas9 score^15^, we selected a highly active sgRNA (id 1) that enables PE3 (Supplementary Table 3, Supplementary Fig. 2a).

We next delivered plasmids encoding the selected pegRNA, the sgRNA, and PE2 (Supplementary Fig. 2b-d) into 5 to 7 week old *Fah*^*mut/mut*^ mice using hydrodynamic injection, after which the mice were treated with NTBC for 7 days (Fig. 3a). After discontinuation of NTBC, all mice that had received PE3 (i.e., PE2, pegRNA, and sgRNA) survived until the end of the experiment (40 days), whereas all mice injected with phosphate-buffered saline (PBS) as a negative control showed substantial weight loss and died before 30 days (Fig. 3b). This extended survival and the prevention of weight loss suggest PE3-induced amelioration of the disease phenotype, which is consistent with the results of previous studies involving genome editing in this mouse model^6-8, 10, 11^.

**Figure 3.**
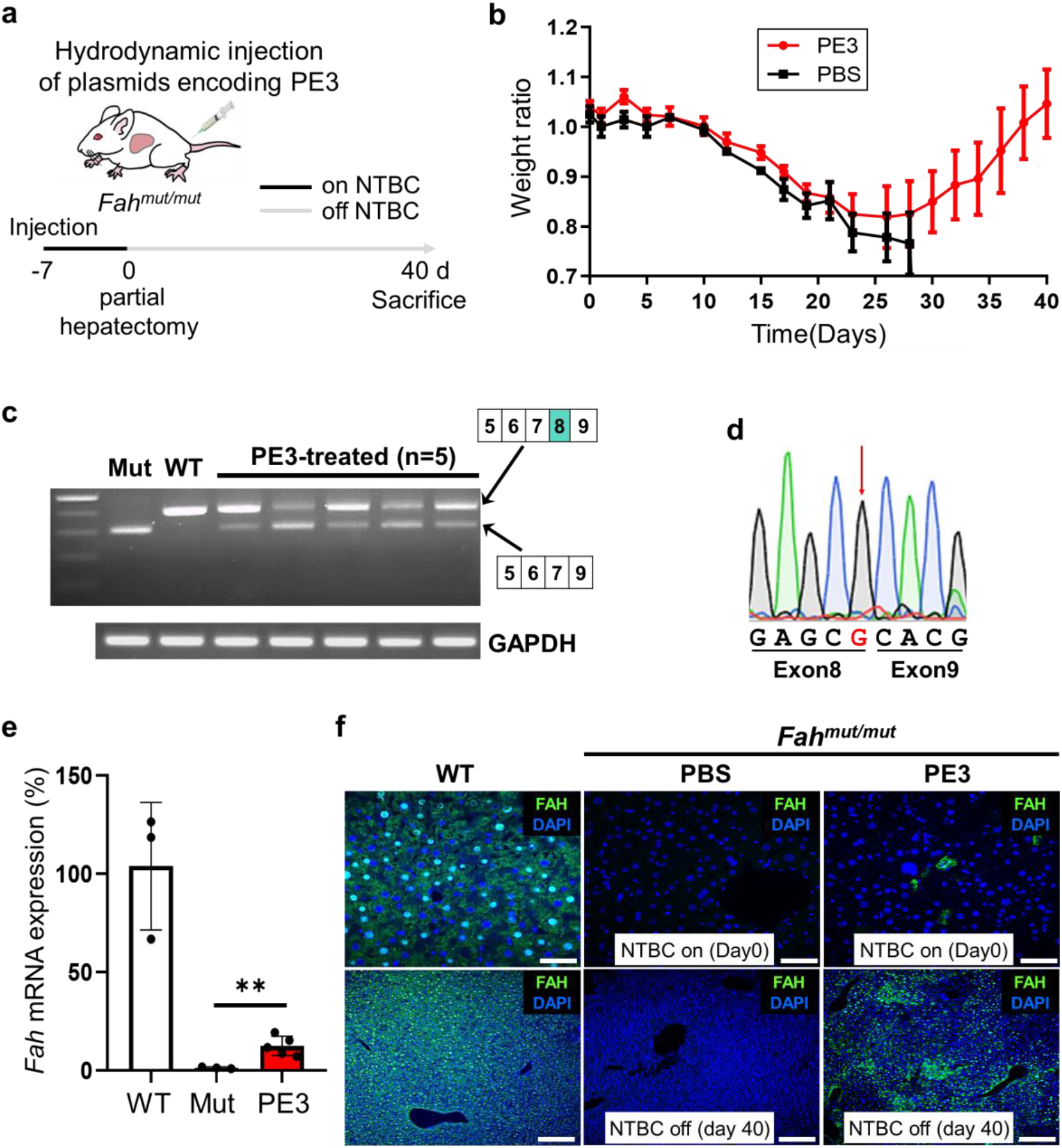
Prime editor 3 corrects the disease mutation and phenotype in *Fah*^*mut/mut*^ mice. (**a**) A schematic representation of the experiments. *Fah*^*mut/mut*^ mice underwent hydrodynamic injection of plasmids encoding prime editor 3 components (i.e., prime editor 2, pegRNA, and sgRNA) and were kept on water containing NTBC for 7 days. The day on which NTBC was withdrawn is defined as day 0. At day 0, a partial hepatectomy was performed to collect liver tissue. At 40 days, the PE3-treated mice were euthanized and analyzed. (**b**) Body weight of *Fah*^*mut/mut*^ mice injected with PE3 or phosphate-buffered saline (PBS, control). Body weights were normalized to the pre-injection weight. The number of mice n = 5 for the PE3 group and n = 3 for the PBS group. Data are mean ± s.e.m. (**c**) Representative RT–PCR using RNA isolated from the liver at 40 days. The primers hybridized to sequences in exons 5 and 9. The wild-type *Fah* (+/+) amplicon is 405 bp in length and the mutant amplicon (which lacks exon 8) is 305 bp. *Gapdh* was used as a control. (**d**) Representative results from Sanger sequencing of the 405-bp RT– PCR band from the PE3-treated mice shown in (c). The red arrow indicates the corrected G nucleotide, which is shown in red type. (**e**) The amount of wild-type *Fah* mRNA in the liver was measured by quantitative RT-PCR using primers that hybridze to sequences in exons 8 and 9. WT, wild-type mice; Mut, *Fah*^*mut/mut*^ mice; PE3, *Fah*^*mut/mut*^ mice injected with plasmids encoding PE3 components. Data are mean ± s.d. The number of mice n = 3 (WT), 3 (Mut), and 5 (PE3). ∗∗p-value = 0.0093. (**f**) Immunofluorescence staining of FAH protein. Scale bars, upper panels, 50 μm; lower panels, 200 μm.

To evaluate whether prime editing rescues exon 8 skipping, at the end of the experimental period (40 days) we conducted reverse-transcription PCR (RT-PCR) using liver mRNA as the template and primers binding exons 5 and 9^11^. A 305-bp PCR amplicon, which indicates exon 8 skipping, was observed for *Fah*^*mut/mut*^ mice, whereas a single 405-bp PCR amplicon, which indicates that exons 5 to 9 are intact, was seen for wild-type mice (Fig. 3c). All five mice injected with PE3 showed both 305- and 405-bp amplicons, suggesting that exon 8 skipping was rescued in a fraction of hepatocytes. Sequencing of the 405-bp amplicon confirmed that the mutant sequence was corrected at the mRNA level (Fig. 3d). Quantitative RT-PCR revealed that the relative average level of exon 8-containing *Fah* mRNA in PE3-treated *Fah*^*mut/mut*^ mice was 12% of that in wild-type mice, whereas such mRNA was not detectable in control *Fah*^*mut/mut*^ mice (Fig. 3e), corroborating that PE3 corrected the exon 8-skipping mutation.

We next quantified the frequency of FAH^+^ cells in the livers of PE3-treated mice. Immunofluorescence staining showed that FAH^+^ cells were present at an average frequency of 0.07% (range, 0.01% to 0.12%) at day 0 (the day NTBC was discontinued) and at an average frequency of 61% (range, 45% to 75%) at day 40 (Fig. 3f, Supplementary Fig. 3). Deep sequencing of liver DNA revealed that the intended edit was not detectable at day 0, which is in line with a previous study of HDR-based genome editing using this mouse model^8^, and was present at an average frequency of 11.5% (range, 6.7% to 18%) at day 40 (Fig. 4a). The reason that the frequency of FAH^+^ cells is higher than the frequency of editing at the DNA level would be because the majority of hepatocytes are polypoid^16^ and because nonparenchymal cell DNA is mixed with that of hepatocytes; similar results were observed in the previous genome editing studies using this mouse model^8, 11^. The observed editing efficiencies are comparable to those obtained with previous approaches using HDR (9.3% at 33 days after the delivery of genome editing components and 30 days after NTBC withdrawal)^8^, microhomology-mediated end joining (5.2% at 37 days after the delivery of genome editing components and 30 days after NTBC withdrawal)^10^, and base editing (9.5% at 38 days after the delivery of genome editing components and 32 days after NTBC withdrawal)^11^ in this mouse model when the genome editing components were delivered using hydrodynamic injections, although direct comparisons are difficult due to the differences in the time points at which the editing efficiencies were analyzed and at which NTBC was discontinued.

**Figure 4.**
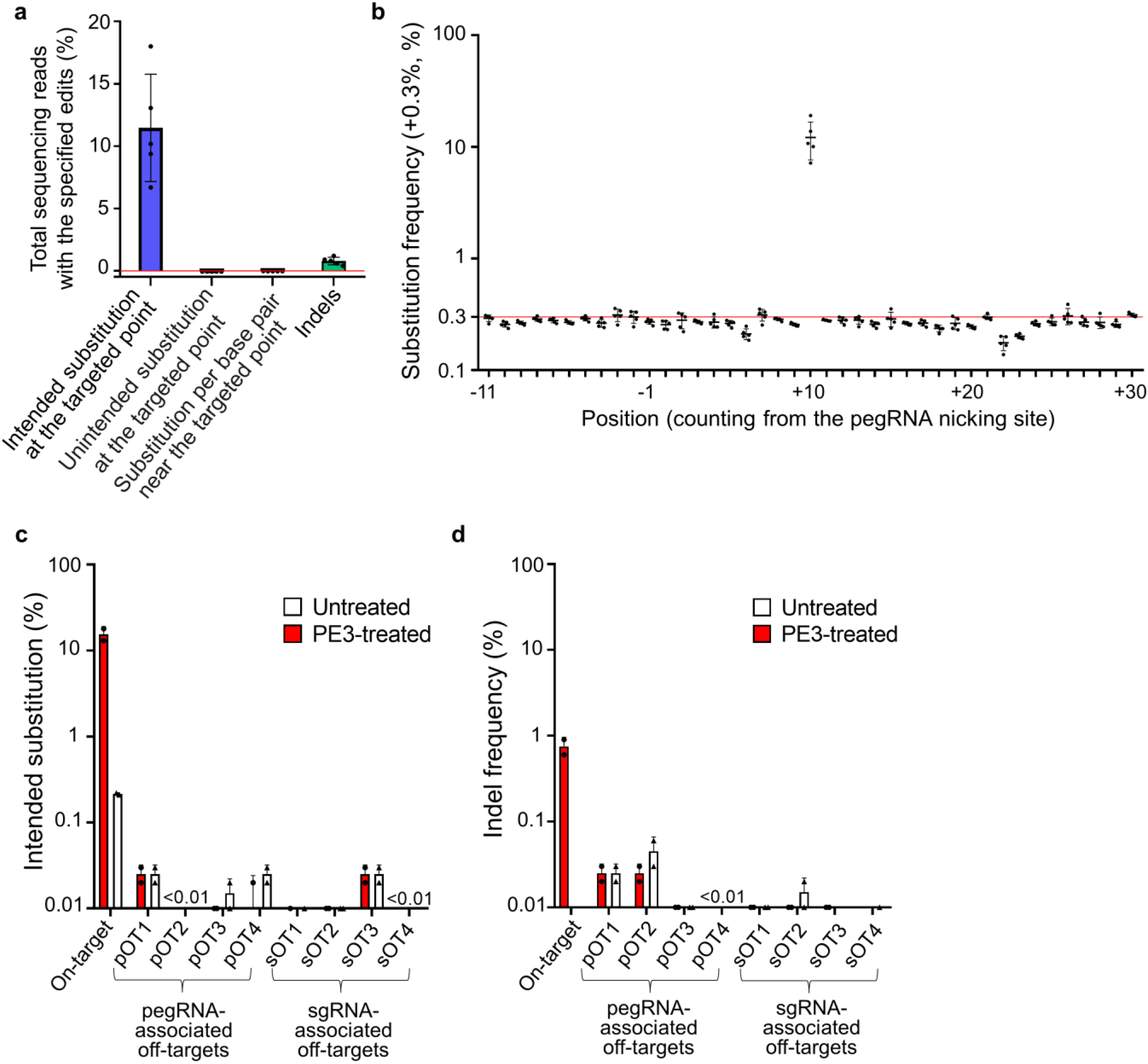
Prime editor 3 corrects the disease-causing mutation in a highly precise manner. (**a**) Frequencies of intended and unintended edits in the livers of PE3-treated mice. The frequencies were normalized by subtracting the average frequency of such editing in the control group injected with PBS to exclude errors originating from PCR amplification and sequencing. Substitutions near the targeted nucleotide were evaluated over a 40-bp range centered on the targeted point. Indels were measured over a 136-bp range centered on the pegRNA nicking site. The red horizontal line represents the position where the normalized frequency = 0. Data are mean ± s.d. The number of mice n = 5. (**b**) Substitution frequencies at each position of the target sequence. The substitution frequency at each position ranging from −20 bp to +20 bp of the targeted position in PE3-treated mice. The frequencies were normalized by subtracting the average edit frequencies in the control group injected with PBS to exclude errors originating from PCR amplification and sequencing. The red horizontal line represents the position where the normalized frequency = 0. Positions are numbered from the pegRNA nicking site. The targeted position is at +10. Data are mean ± s.d. The number of mice n = 5. (**c, d**) Frequences of the intended substitution (c) and indels (d) at the four top-ranking predicted off-target sites for the used pegRNA (pOT1, .., pOT4) and those for the used sgRNA (sOT1, …, sOT4) in PE3-treated liver tissues. Genomic DNA isolated from the livers of *Fah*^*mut/mut*^ mice without PE3 treatment was used as the negative control (Untreated). The number of mice n = 2.

### Prime editor 3 corrects the disease-causing mutation in a highly precise manner

We next determined whether PE3 induced any unintended editing including indels in or near the target sequence in the mouse liver. Deep sequencing revealed that unintended substitutions at or near the targeted nucleotide were not detected in any of the mice and that the level of indels ranged from only 0.4% to 1.2% (on average, 0.78%) (Fig. 4a, b). In the case of the Cas9-directed HDR approach, the frequency of indels observed at the target site was 26%, which is 33-fold higher than the level induced by PE3 in this study. When adenosine base editor was used, the frequency of unintended substitutions observed near the target nucleotide was 1.9%^11^. Thus, these data suggest that prime editing can be more precise than the other approaches in mouse somatic cells.

To quantify the off-target effects of PE3, we identified the four top-ranking potential off-target sites of the used pegRNA and the four top-ranking potential off-target sites of the used sgRNA using CRISPOR^17^ (Supplementary Table 4). Deep sequencing revealed no off-target effects, including intended substitutions or indels, at any of the eight sites (Fig. 4c, d). This highly specific editing by prime editor 3 is in line with recent results showing that off-target effects were not detectable when PE3 was used in mouse embryos^5^ or human organoids^3^ and is also compatible with the low frequency of off-target effects of PE3 in cultured mammalian cells^1, 18^.

### Prime editor 2 corrects the disease mutation and phenotype in *Fah*^*mut/mut*^ mice

We also performed similar experiments using PE2, which does not require an sgRNA, instead of PE3 (Supplementary Fig. 4a). *Fah*^*mut/mut*^ mice treated with PE2 and the pegRNA used for the PE3 experiments described above survived until the end of the experiment (60 days after the initial NTBC withdrawal), whereas the control *Fah*^*mut/mut*^ mice all died within 30 days (Supplementary Fig. 4b). Quantitative RT-PCR showed that the expression level of *Fah* mRNA containing intact exon 8 in PE2-treated mutant mice was on average 6.9% that in wild-type mice, but that the expression in untreated mutants was undetectable (Supplementary Fig. 4c). Immunofluorescence staining showed that an average of 33% of liver cells from PE2-treated mutant mice were FAH^+^ at 60 days (Supplementary Fig. 4d).

### Prime editor 2 corrects the mutation without any detectable unintended substitutions, indels, bystander effects, or off-target effects

We analyzed genomic DNA isolated from the PE2-injected mice at the end of the experiments (60 days after the initial NTBC withdrawal) by deep sequencing. We found that the intended edit was present in an average of 4.0% of the total sequencing reads (Fig. 5a). Any unintended edits including indels, unintended substitutions at the targeted nucleotide, and bystander nucleotide edits were undetectable (Fig. 5a, b). Furthermore, when we evaluated the four potential off-target sites of the pegRNA predicted by CRISPOR^17^ (Supplementary Table 4), no off-target effects, including substitution mutations or indels, were detected, either (Fig. 5c, d). Taken together, these results suggest that PE2-mediated prime editing in mice can correct the disease-causing mutation, in a highly precise and specific manner, which could not have been achieved using other previous genome editing approaches based on either a CRISPR nuclease or a base editor.

**Figure 5.**
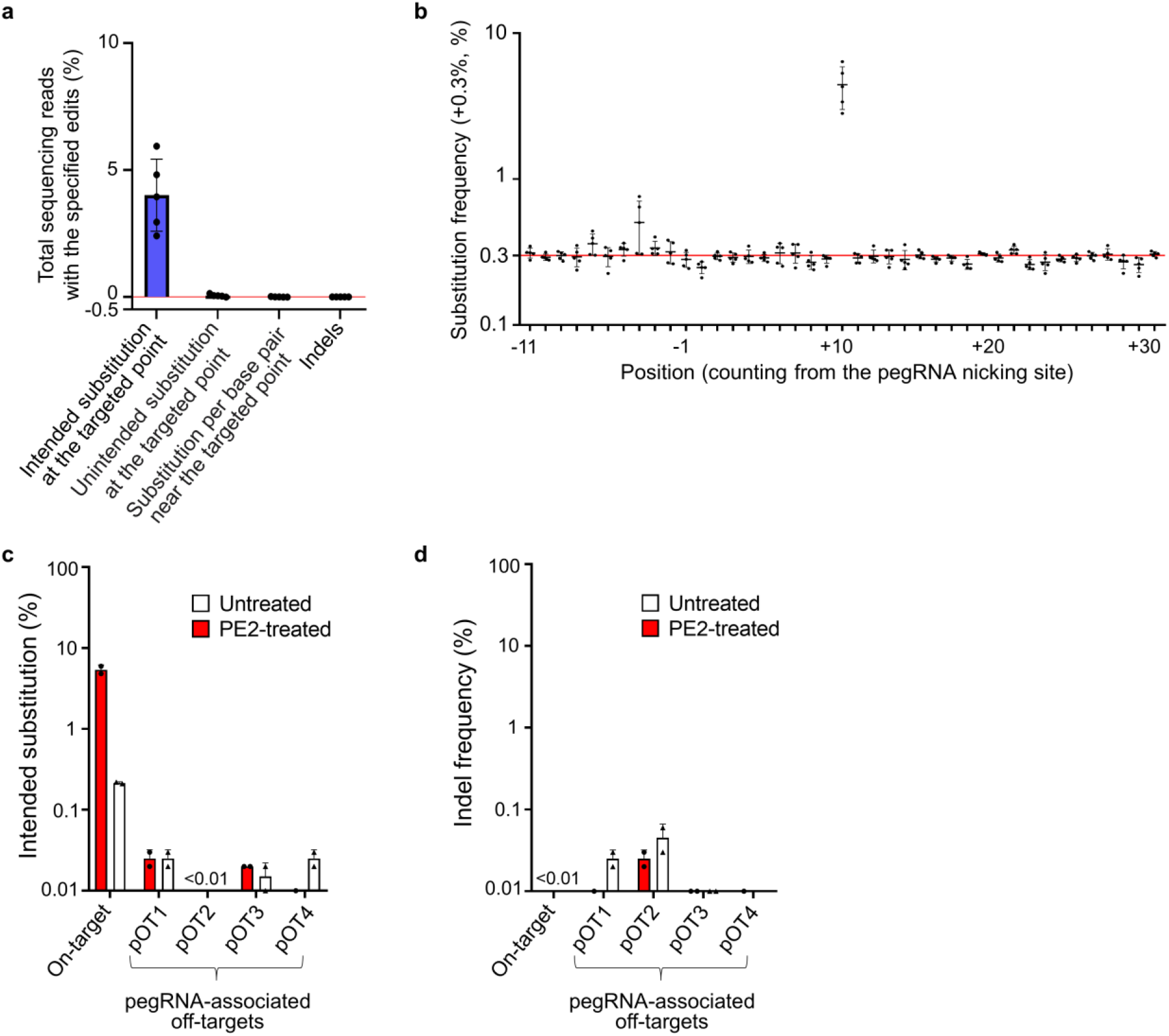
Prime editor 2 corrects the disease-causing mutation without any detectable unintended substitutions, indels, bystander effects, or off-target effects. (**a**) Frequencies of intended and unintended edits in the livers of PE2-treated mice. The frequencies were normalized by subtracting the average frequency of such editing in the control group injected with PBS to exclude errors originating from PCR amplification and sequencing. Substitutions near the targeted nucleotide were evaluated over a 40-bp range centered on the targeted point. Indels were measured over a 136-bp range centered on the pegRNA nicking site. The red horizontal line represents the position where the normalized frequency = 0. Data are mean ± s.d. The number of mice n = 5. (**b**) Substitution frequencies at each position of the target sequence. The substitution frequency at each position ranging from −20 bp to +20 bp of the targeted position in PE2-treated mice. The frequencies were normalized by subtracting the average edit frequencies in the control group injected with PBS to exclude errors originating from PCR amplification and sequencing. The red horizontal line represents the position where the normalized frequency = 0. Positions are numbered from the pegRNA nicking site. The targeted position is at +10. Data are mean ± s.d. The number of mice n = 5. (**c, d**) Frequencies of the intended substitution (c) and indels (d) at the four top-ranking predicted off-target sites (pOT1, .., pOT4) for the used pegRNA in PE2-treated liver tissues. Genomic DNA isolated from the livers of *Fah*^*mut/mut*^ mice without PE2 treatment was used as the negative control (Untreated). The number of mice n = 2.

### AAV-mediated prime editing in the retina and RPE of wild-type mice

We next investigated whether prime editing could be achieved in another adult tissue using a different delivery method. Given that there are several genetic retinal diseases that could potentially be treated using therapeutic genome editing^19^, we chose to test in vivo prime editing in the retina of adult mice. We also chose adeno-associated virus (AAV) for the delivery of PE2-, pegRNA-, and sgRNA-encoding sequences, because AAV has been used to efficiently deliver sequences encoding other genome editing tools including engineered nucleases and base editors. Given that the coding sequence of PE2 is 6,273 bp, which is longer than the cargo size limit of AAV, we used trans-splicing AAV (tsAAV) vector^20-22^ (serotype 8) (Fig. 6a). We chose to target the *ATP7B* gene, because editing this gene is not expected to affect the viability or function of retinal cells. We designed four pegRNAs that would induce a G-to-A substitution and chose the one associated with the highest activity (8.3%) in cultured neuro2A cells (Fig. 6b, 6c). An sgRNA that was predicted to be highly active was selected using DeepSpCas9^15^ (Supplementary Table 5).

**Figure 6.**
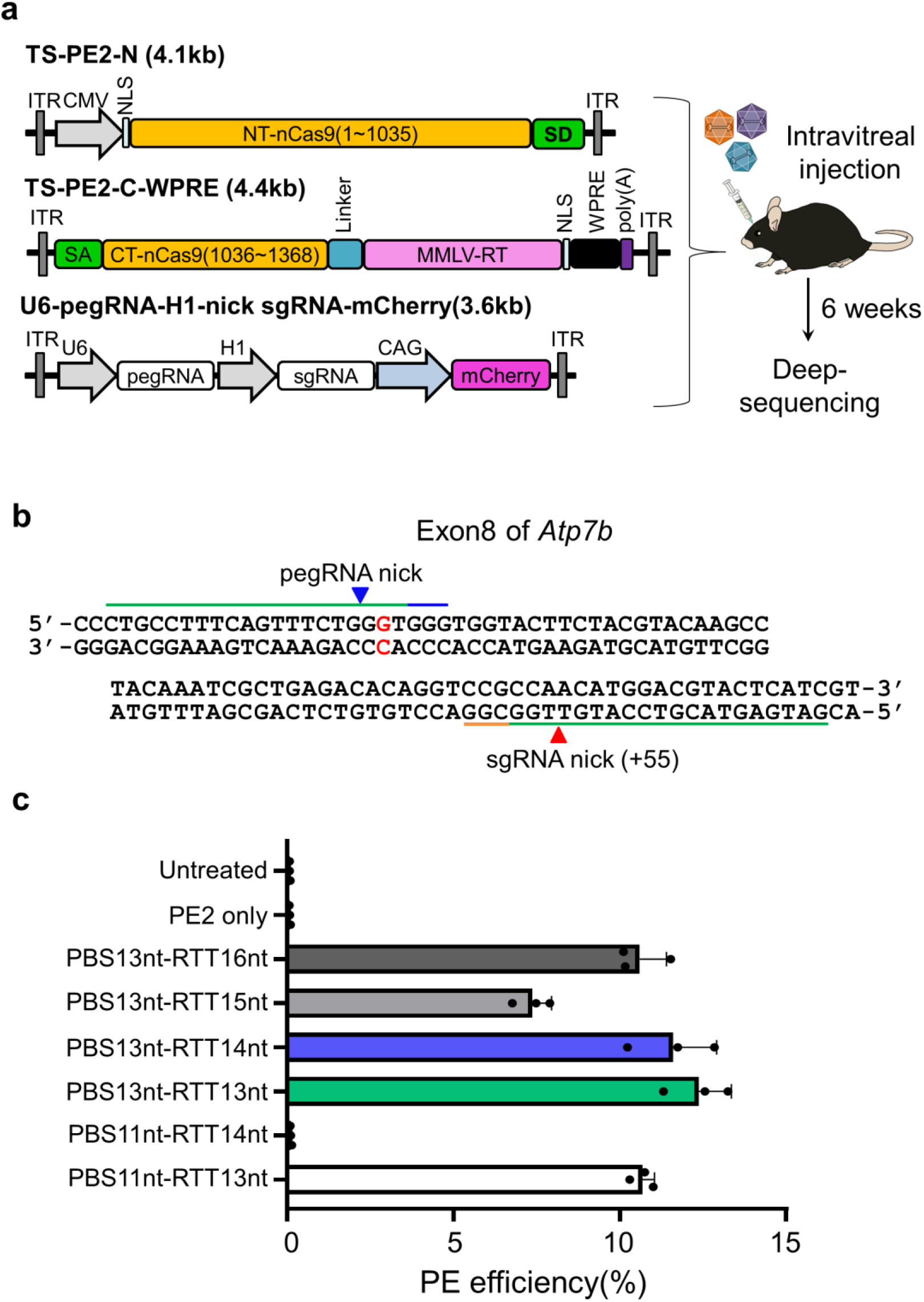
AAV-mediated prime editing in the retina and RPE of wild-type mice. (**a**) A schematic representation of experiments together with maps of the AAV vectors used in the experiment. The lengths of the sequences between the two ITRs in each vector are shown in parentheses. Two transplicing PE2-expressing AAVs (TS-PE2-N and TS-PE2-C-WPRE) were delivered together with an AAV expressing pegRNA and sgRNA into the retina and RPE via intravitreal injection. The retina and RPE cells were harvested for deep sequencing at 6 weeks after the injection. SD, splicing donor; SA, splicing acceptor; NLS, nuclear localization sequence; ITR, inverted terminal repeat. (**b**) The target and neighboring sequence in exon 8 of *Atp7b*. The green lines represent the pegRNA and sgRNA spacers. The PAMs of the pegRNA and sgRNA target sequences are highlighted in blue and orange, respectively. (b) Editing frequencies of the six pegRNAs measured in N2A cells. Six pegRNAs targeting *Atp7b* exon 8 were designed to induce a G>A mutation. The six pegRNAs differed in the lengths of their primer binding sites and RT templates, but shared a common target sequence. The pegRNA harboring a 13-nt PBS and 13-nt RT template was used for in vivo delivery.

### AAV-mediated prime editing in the retina and RPE did not induce any detectable unintended substitutions, indels, bystander effects, or off-target effects

We delivered two tsAAV vectors, one encoding the N-terminal half of PE2 and the other encoding the C-terminal half, together with an AAV vector encoding the *ATP7B*-targeting pegRNA and sgRNA, into the mouse retina and retinal pigment epithelium (RPE) using intravitreal injection. Six weeks after the injection, the mice were euthanized, and the RPE and retina were harvested and analyzed (Fig. 6a). Deep sequencing showed that the average editing efficiency was 1.82% and 1.87% in the RPE and retina, respectively, without detectable indels or unintended substitutions at or nearby the targeted nucleotide (Fig. 7a-c, Supplementary Figs. 5 and 6). These data suggest that AAV-mediated PE3 delivery can induce precise genome editing in the retina and RPE of adult mice. Given that a frequency of even 1.17% of the desired edit can lead to a substantial improvement in retina function in a mouse model of a human genetic eye disease^23^, this level of PE3-induced genome editing could be useful for therapeutic genome editing.

**Figure 7.**
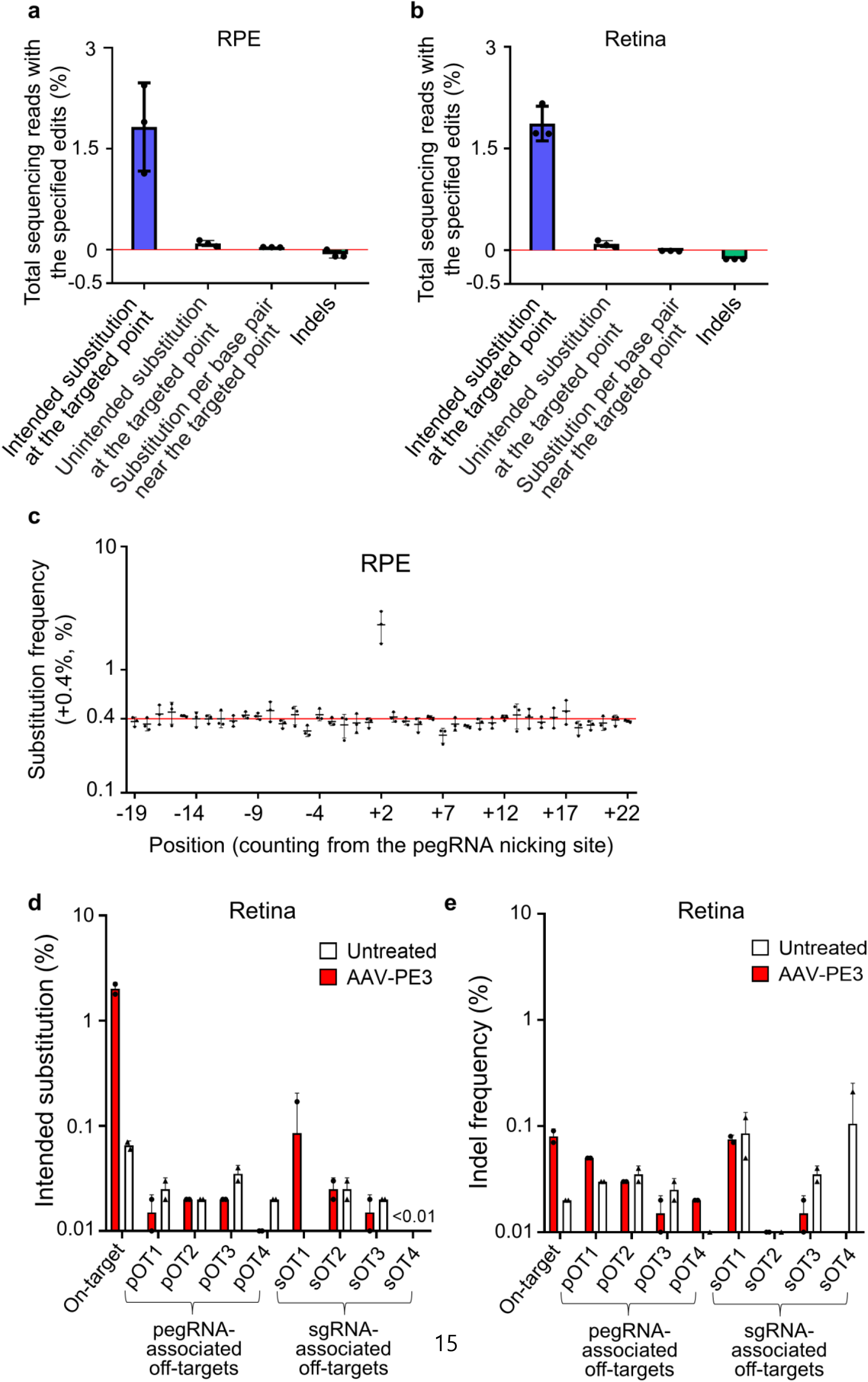
AAV-mediated prime editing in the retina and RPE did not induce any detectable unintended substitutions, indels, bystander effects, or off-target effects. (**a**) Frequencies of intended and unintended edits in the RPE (a) and retina (b) of PE3-treated mice. The frequencies were normalized by subtracting the average frequencies of such edits in the control group injected with PBS to exclude errors originating from PCR amplification and sequencing. Substitutions near the targeted nucleotide were evaluated over a 40-bp range centered on the targeted point. Indels were measured over a 136-bp range centered on the pegRNA nicking site. The red horizontal line represents the position where the normalized frequency = 0. Data are mean ± s.d. The number of mice n = 3. (**c**) The substitution frequency at each position ranging from −20 bp to +20 bp of the targeted position in the RPE treated with PE3. The frequencies were normalized by subtracting the background substitution frequencies in controls that were not treated with PE3. The red horizontal line represents the position where the normalized frequency = 0. Positions are numbered from the pegRNA nicking site. The targeted position is at +2. The targeted position is at +2. Data are mean ± s.d. The number of mice n = 3. (**d, e**) Frequencies of the intended substitution (d) and indels at the four top-ranking predicted off-target sites for the used pegRNA (pOT1, .., pOT4) and those for the used sgRNA (sOT1, …, sOT4) in PE3-treated retinas. Genomic DNA isolated from the retinas of mice without PE3 treatment was used as the negative control (Untreated). The number of mice n = 2.

We next found the four top-ranking potential off-target sites of the pegRNA and those of the sgRNA using CRISPOR^17^(Supplementary Table 4). Deep sequencing at those eight sites revealed no detectable off-target substitutions or indels (Fig. 7d, e), corroborating that prime editing is highly specific.

## Discussion

Our results demonstrate that prime editing can generate intended edits in a highly precise manner in the liver and eye of adult mice when components are delivered using hydrodynamic injection or AAV-mediated methods. We also showed that replacing the SpCas9 domain of PE2 with a PAM variant such as SpCas9-NG can expand the list of target sequences for prime editors.

Cells containing disease-causing mutations are often not readily available, which makes it difficult to evaluate the efficiencies of genome editing tools including prime editors at the mutant target sequence. Furthermore, hundreds or thousands of pegRNAs can be designed to induce an intended edit at a target sequence. Determining pegRNA activities using lentiviral libraries of paired pegRNA and target sequences as we describe above would be useful especially when mutant sequence-containing cell lines are not available or when a large number of pegRNAs are evaluated.

When we compared PE2- and PE3-based correction of the disease-causing mutation in *Fah*^*mut/mut*^ mice using hydrodynamic injections with those based on the hydrodynamic injections of Cas9 nuclease^8, 10^ or adenosine base editor^8^, the efficiencies were overall comparable although exact comparisons are difficult, at least in part, due to differences in the time points used for analysis (Supplementary Table 6). However, the most striking difference is the precision of the editing. Cas9 nuclease induced a substantial frequency of indels (26%) at the target sites and a detectable, albeit lower than 0.3%, frequency of indels at off-target sites^8^, whereas adenosine base editor induced significant bystander effects (1.9%). However, PE3, but not PE2, induced only a low level of indels (on average 0.78%) and bystander or off-target effects were not observed for either PE2 or PE3. The level of precision in genome editing we observed for PE2 in particular has not been achieved using any of the previous methods of genome editing in this mouse model.

This high precision of prime editing in somatic cells of mice raises the possibility that prime editing could be used for genome editing in human patients. We envision that in vivo prime editing, together with previously reported base editing and engineered nuclease-based approaches, will be a promising tactic for genome editing therapy for genetic diseases.

## Methods

### Construction of plasmid vectors

To prepare pLenti-NG-PE2-BSD, the Lenti_Split-BE4-N-Blast plasmid^24^ was digested with restriction enzymes AgeI and BamHI (New England Biolabs (hereafter, for brevity, NEB)) and a MEGAquick-spin total fragment DNA purification kit (iNtRON Biotechnology) was used to gel purify the linearized plasmid. Fragments of PE2-encoding sequence (Addgene #132775) and NG-Cas9-encoding sequence (Addgene #124163) were amplified by PCR using Phusion Polymerase (NEB). The amplicons and the linearized plasmid were assembled using an NEBuilder HiFi DNA assembly kit (NEB). The assembled plasmid was named pLenti-NG-PE2-BSD.

To prepare px601-PE2, the PE2-encoding sequence (Addgene #132775) and the WPRE sequence (Addgene #52962) were amplified by PCR and cloned into the pX601 plasmid using an NEBuilder HiFi DNA assembly kit (NEB).

To construct px552-pegRNA-CAG-mCherry, gRNA scaffold sequence (Addgene #104174) and CAG-mCherry (Addgene #108685) fragments were PCR-amplified and cloned into the px552 plasmid (Addgene #60958) to make px552-U6-CAG-mCherry. Next, pegRNA sequences were synthesized and cloned into the linearlized px552-U6-CAG-mCherry plasmid, which had been digested with SapI and SpeI (NEB), using an NEBuilder HiFi DNA assembly kit (NEB). These manipulations generated px552-pegRNA-CAG-mCherry.

An sgRNA-expressing vector (Addgene #104174) was digested with BsmBI. sgRNA oligomers were annealed, phosphorylated with T4 PNK, and ligated with the linearized vector to construct gN19-nicking sgRNA.

### Construction of a plasmid library of pegRNA and target sequence pairs

Oligonucleotides were designed and the library of pegRNA-encoding and target sequence pairs was generated as previously described^2^. Briefly, an oligonucleotide pool containing 435 pairs of pegRNA-encoding sequences and target sequences was synthesized by Twist Bioscience (San Francisco, CA). Each oligonucleotide included the following elements: a 19-nt guide sequence, BsmBI restriction site #1, a 15-nt barcode stuffer sequence, BsmBI restriction site #2, the RT template sequence, the primer binding site, a poly T sequence, an 18-nt barcode sequence (identification barcode), and a corresponding 43∼47-nt wide target sequence that included a PAM and an RT template binding region. Oligonucleotides that contained other, unintended BsmBI sites were excluded. The barcode stuffer was later excised by digestion with BsmBI; the identification barcode (located upstream of the target sequence) allowed individual pegRNA and target sequence pairs to be identified after deep sequencing.

The plasmid library containing pairs of pegRNA-encoding and target sequences was prepared using a two-step cloning process^2^. This method effectively prevents uncoupling between paired guide RNA and target sequences during PCR amplification of oligonucleotides^25^.

### Production of lentivirus

8 × 10^6^ HEK293T cells were seeded on 150-mm cell culture dishes containing Dulbecco’s Modified Eagle Medium (DMEM). After incubation for 16 hours, the DMEM was exchanged with fresh medium containing 25 μM chloroquine diphosphate; the cells were then incubated for 4 more hours. The plasmid library, psPAX2 (Addgene #12260), and pMD2.G (Addgene #12259) were mixed to yield a total of 40 μg of the plasmid mixture. HEK293T cells were then transfected with this mixture using polyethyleneimine. Fifteen hours later, cultures were refreshed with maintaining medium. At 48 hours after transfection, the supernatant (containing lentivirus) was collected, filtered through a Millex-HV 0.45-μm low protein-binding membrane (Millipore), and aliquoted. Serial dilutions of a viral aliquot were prepared and transduced into HEK293T cells so that the virus titer could be determined. Untransduced cells and cells treated with the serially diluted virus were both cultured in the presence of 2 µg/ml puromycin (Invitrogen). The lentiviral titer was quantified by counting the number of living cells at the time when all of the untransduced cells died as previously described^26^.

### Cell library generation and PE2 delivery

The cell library was generated as previously described^2^. Briefly, HEK293T cells were seeded on eighteen 150-mm dishes (at a density of 1.2 × 10^7^ cells per dish) and incubated overnight. The lentiviral library was transduced into the cells at a multiplicity of infection of 0.3 to achieve >500× coverage relative to the initial number of oligonucleotides. After incubation of the cells overnight, untransduced cells were removed by maintaining the cultures in 2 µg/ml puromycin for the next 5 days. The cell library was maintained at a count of at least 7.2 × 10^7^ cells for the entire study to preserve library diversity. Next, a total of 7.2 x 10^7^ cells (from six 150-mm culture dishes, each with 1.2 x 10^7^ cells) were transfected with the pLenti-NG-PE2-BSD plasmid (80 µg per dish) using 80 µl Lipofectamine 2000 (Thermo Fisher Scientific) according to the manufacturer’s instructions. Six hours later, the culture medium was replaced with DMEM supplemented with 10% fetal bovine serum and 20 µg/ml blasticidin S (InvivoGen). Five days later, the cells were harvested for genomic DNA extraction.

### Deep sequencing

Genomic DNA was extracted from harvested cells with a Wizard genomic DNA purification kit (Promega) and from mouse tissue (liver, retina, RPE) using a DNeasy Blood & Tissue kit (Qiagen). Target sequences were PCR-amplified using 2X Taq PCR Smart mix (SolGent). The PCR primers used for the experiment are shown in Supplementary Table 7.

In preparation for evaluating the activities of the 435 pegRNAs in a high-throughput manner, an initial PCR included a total of 350 μg genomic DNA; assuming 10 μg genomic DNA per 10^6^ cells, coverage would be more than 1000× over the library. 3.64 μg of genomic DNA per reaction (with a total of 96 reactions) was amplified using primers that included Illumina adaptor sequences, after which the products were pooled and gel-purified with a MEGAquick-spin total fragment DNA purification kit (iNtRON Biotechnology). Next, 100 ng purified DNA was amplified by PCR using primers that included barcode sequences. After gel purification, the amplicons were analyzed using the NovaSeq platform (Illumina, San Diego, CA).

For evaluation of prime editing at endogenous sites, the first PCR for the amplification of the target sequence was performed in a 50-µL reaction volume that contained 3 μg of the initial genomic DNA template for liver samples and 500 ng of DNA template for retina and RPE samples. The second PCR to attach the Illumina barcode sequences was then performed using 50 ng of the purified product from the first PCR in a 30-µl reaction volume. The resulting amplicons were sequenced after gel purification using MiniSeq (Illumina, San Diego, CA) according to the manufacturer’s protocols.

### Analysis of prime editing efficiencies

Previously reported Python scripts^2^ were used to analyze the high-throughput results. Each pegRNA and target sequence pair was identified using a 22-nt sequence (the 18-nt barcode and 4-nt sequence located upstream of the barcode). Reads that included the specified edits but not unintended mutations within the wide target sequence were considered to contain NG-PE2-induced mutations. To exclude the background mutations generated during the oligo synthesis and PCR amplification steps, we subtracted the background prime editing frequencies determined in the cell library that had not been treated with NG-PE2 from the observed prime editing frequencies as previously described^2^. pegRNA and target sequence pairs with deep sequencing read counts below 100 and those with background prime editing frequencies above 5% were removed from the analyses as previously described^27^.

To quantify the prime editing frequencies at endogenous sites, amplicon sequences were aligned to reference sequences using Cas-analyzer^28^. The frequencies of intended and unintended edits, including substitutions and indels, were calculated as the percentage of (number of reads with the edit/number of total reads). The frequencies of such edits in the experimental groups were normalized by subtracting the average frequency of such edits in the control groups to exclude errors originating from PCR amplification and sequencing.

### Generation of mutant *Fah* target sequence-containing HEK293T cells

The target region of the *Fah* gene was PCR-amplified from genomic DNA from the *Fah*^*mut/mut*^ mouse and cloned into a lentivirus shuttle vector derived from a hygromycin reporter plasmid^29^. Next, lentivirus was produced from the vector and transduced into HEK293Tcells. After incubation of the cells overnight, untransduced cells were removed by supplementing the culture medium with 2 μg/ml hygromycin for the next 5 days.

### Analysis of off-target effects

Potential off-target sites were computationally identified using CRISPOR^17^. The four top-ranking predicted off-target sites of the used pegRNA and those of the used sgRNA were analyzed by targeted deep sequencing. The sequences of the off-target sites are provided in Supplementary Table 4. The primers used for deep sequencing are shown in Supplementary Table 7. Cas-analyzer^28^ was used to analyze indel frequencies at off-target sites and the frequencies of the intended edit were analyzed using the same method used to analyze frequencies at the on-target sites described above.

### *Fah*^*mut/mut*^ mice and hydrodynamic injection

All animal study protocols were approved by the Institutional Animal Care and Use Committee (IACUC) of Yonsei University Health System (Seoul, Korea). *Fah*^*mut/mut*^ mice^6^ were provided with water containing 10 mg/L NTBC unless specified. Plasmids for hydrodynamic tail-vein injection were prepared using an EndoFree Plasmid Maxi kit (Qiagen). Plasmids encoding PE2 (127 μg) and pegRNA (73 μg), which had been suspended in 2 ml normal saline, were injected into the tail vein of 5 to 7 week old *Fah*^*mut/mut*^ mice within 5-7 seconds (30 μg of sgRNA plasmid was added to the normal saline only for the PE3-treated group). For measuring the initial correction efficiency, *Fah*^*mut/mut*^ mice were kept on water containing NTBC for 7 days after injection with plasmid DNA; at that time, NTBC was withdrawn and a partial hepatectomy was performed. PE2-treated mice were sacrificed at 60 days after NTBC withdrawal for histology and DNA and RNA analysis. PE3-treated mice were sacrificed at 40 days after NTBC withdrawal.

### AAV vector construction and virus production

To prepare the trans-splicing PE2 plasmid, the sequence encoding the PE2 N terminus (PE2-N term) (Addgene #132775) and a splicing donor (SD) (Addgene#112734) were PCR-amplified using primers listed in Supplementary Table 7 and cloned into the px601 plasmid using an NEBuilder HiFi DNA assembly kit (NEB). Similarly, a splicing acceptor (SA) (Addgene#112876), the sequence encoding the PE2 C terminus (PE2-C term) (Addgene #132775), and the WPRE sequence (Addgene #52962) were PCR-amplified and cloned into the pX601 plasmid using an NEBuilder HiFi DNA assembly kit (NEB).

To construct U6-pegRNA-H1-nick sgRNA-mCherry, pegRNA-encoding, sgRNA-encoding, and H1 promoter sequences (Addgene#61089) were either synthesized or PCR-amplified and then cloned into the px552-U6-CAG-mCherry plasmid using an NEBuilder HiFi DNA assembly kit (NEB). The AAV vectors were then sent to VectorBuilder Inc (Chicago, Illinois, USA) for packaging and ultra-purification to obtain the high-grade AAVs (serotype 8) required for an in vivo study.

### Intravitreal injection of AAV

Mouse eye-related studies were performed following the guidelines of the Association for Research in Vision and Ophthalmology statement for the use of animals in ophthalmic and vision research and were approved by the IACUCs of both Seoul National University and Seoul National University Hospital. C57BL/6 mice, obtained from Central Laboratory Animal, were maintained under a 12-hour dark/light cycle. Before AAV injection, deep anesthesia was induced by tiletamine plus zolazepam (Zoletil 50, Virbac; 30 mg/kg) and xylazine (Rompun, Bayer; 10 mg/kg). Then, 4.9 × 10^10^ vg of AAV8-TS-PE2-N term, 4.9 × 10^10^ vg of AAV8-TS-PE2-C term, and 2.4 × 10^10^ vg of AAV8-Atp7b pegRNA-sgRNA in PBS were mixed in a total volume of 1.5 μl and injected into the vitreous cavity of the mouse eye. A customized Nanofil syringe with a blunt 33-gauge needle (World Precision Instrument) was used for the injection, which was performed using an operating microscope (Leica).

### Immunofluorescence

Mice were euthanized by carbon dioxide asphyxiation. Livers were fixed in 4% paraformaldehyde overnight at 4°C, followed by immersion in 6% sucrose overnight and immersion in 30% sucrose the next day. The livers were embedded in OCT compound (Leica, Wetzlar, Germany), cryosectioned at 16 μm, and stained with hematoxylin and eosin (H&E) for pathology studies. The liver cryostat sections were washed three times with 1X PBS and incubated in blocking buffer for 1 hour at room temperature. The sections were then incubated with primary anti-Fah antibody (1:200, Abcam, Cambridge, UK) at 4°C overnight. After washing with 1X PBS, the sections were incubated with Alexa Fluor® 488 goat anti-rabbit secondary antibody (1:400, Invitrogen, Carlsbad, CA, USA) for 1 hour at room temperature. Cells were mounted on glass slides and nuclei were visualized using fluorescent mounting medium containing 4’,6-diamidino-2-phenylindole (DAPI; Vector, Burlingame, CA). Images were captured using a confocal microscope (LSM700, Zeiss, Gottingen, Germany).

Quantification of FAH^+^ hepatocytes in the liver was performed for 3-5 mice per group, from >4 liver regions per mouse, with ZEN Imaging software (Blue edition, Zeiss).

### Gene expression analysis by quantitative RT-PCR

Total RNA was purified using TRIzol (Invitrogen) and reverse-transcribed using AccuPower RT PreMix (Bioneer, Korea). Quantitative RT-PCR was performed using SYBR Green (Applied Biosystems, Carlsbad, CA); the primers are listed in Supplementary Table 7. Gene expression levels were normalized to *Gapdh* levels. All experiments were performed in triplicate.

### Statistics and reproducibility

P values were determined by Student’s t-test using GraphPad Prism8. To determine Pearson and Spearman correlation coefficients, we used Microsoft Excel (version 16.0, Microsoft Corporation). For high-throughput evaluation of pegRNA efficiencies, we combined the data from two replicates independently transfected by two different experimentalists.

### Data availability

The deep sequencing data from this study have been submitted to the NCBI Sequence Read Archive (SRA; https://www.ncbi.nlm.nih.gov/sra/) under accession number SRR12778000. The datasets used in this study are provided as Supplementary Table 2.

## Author contributions

H.J. and J.H.S. performed most of the experiments. D.H.J. and J.H.K. conducted the eye-related experiments. G.Y. and R. G. performed the high-throughput evaluation of PE2 efficiencies. J.H.S. and S.-R.C. conducted immunofluorescence staining for FAH and histologic evaluations. H.J. and H.H.K. analyzed the data and wrote the manuscript.

## Acknowledgments

We would like to thank Sehyuk Kwon and Jinman Park for helping with computational analyses. We are grateful to Seonmi Park and Younghye Kim for assisting with the experiments. This work was supported in part by the National Research Foundation of Korea (grants 2017R1A2B3004198 (H.H.K.), 2017M3A9B4062401 (H.H.K. and J.H.K.), 2020R1C1C1003284 (H.K.K), and 2018R1A5A2025079 (H.H.K)), Brain Korea 21 Plus Project (Yonsei University College of Medicine), and the Korean Health Technology R&D Project, Ministry of Health and Welfare, Republic of Korea (grants HI17C0676 (H.H.K.) and HI16C1012 (H.H.K. and S.-R.C.)), and Creative Materials Discovery Program through the National Research Foundation of Korea (2018M3D1A1058826 (J.H.K.)).

## Notes

### Competing Interest Statement

Yonsei University has filed a patent application based on this work, in which H.J. and H.H.K. are listed as inventors.

