## Supplementary Material for "Prime editing enables precise genome editing in mouse liver and retina"

Contents:

Supplementary Figures 1 – 6

Supplementary Tables 1 – 7

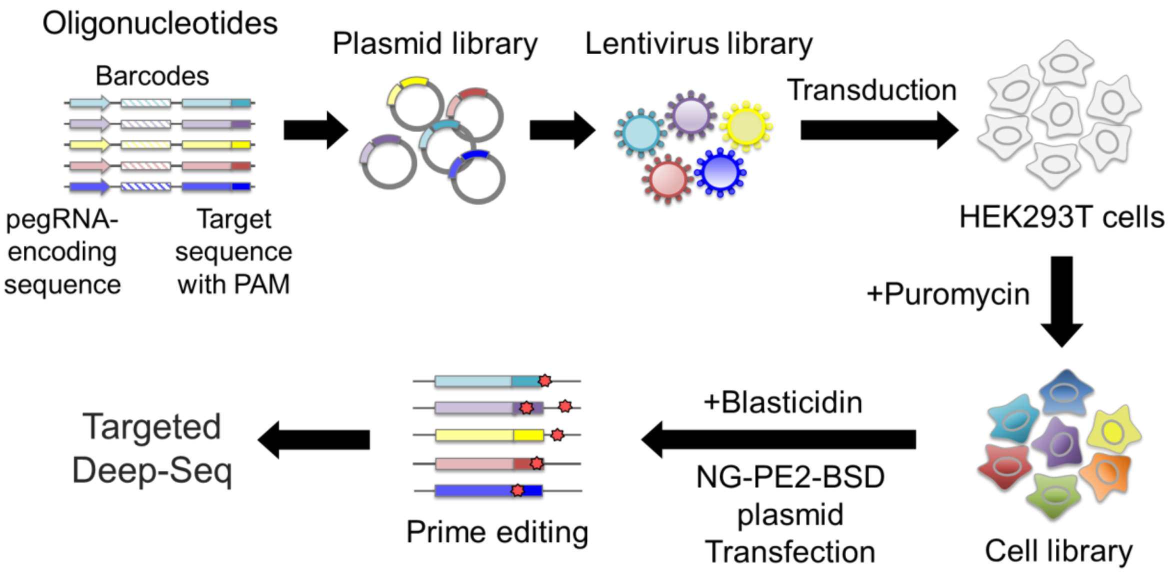


**Supplementary Figure 1.** Schematic representation of the high-throughput evaluation of pegRNA activities. A lentiviral plasmid library was prepared from a pool of oligonucleotides that contained pairs of pegRNA-encoding sequences and corresponding target sequences. Next, HEK293T cells were transduced with lentivirus generated from the plasmid library to construct a cell library and untransduced cells were removed by puromycin selection. This cell library was then transfected with a plasmid encoding NG-PE2, and untransfected cells were removed by blasticidin selection. Five days after the transfection, genomic DNA was isolated from the cells, PCR-amplified, and subjected to deep sequencing to determine prime editing efficiencies.

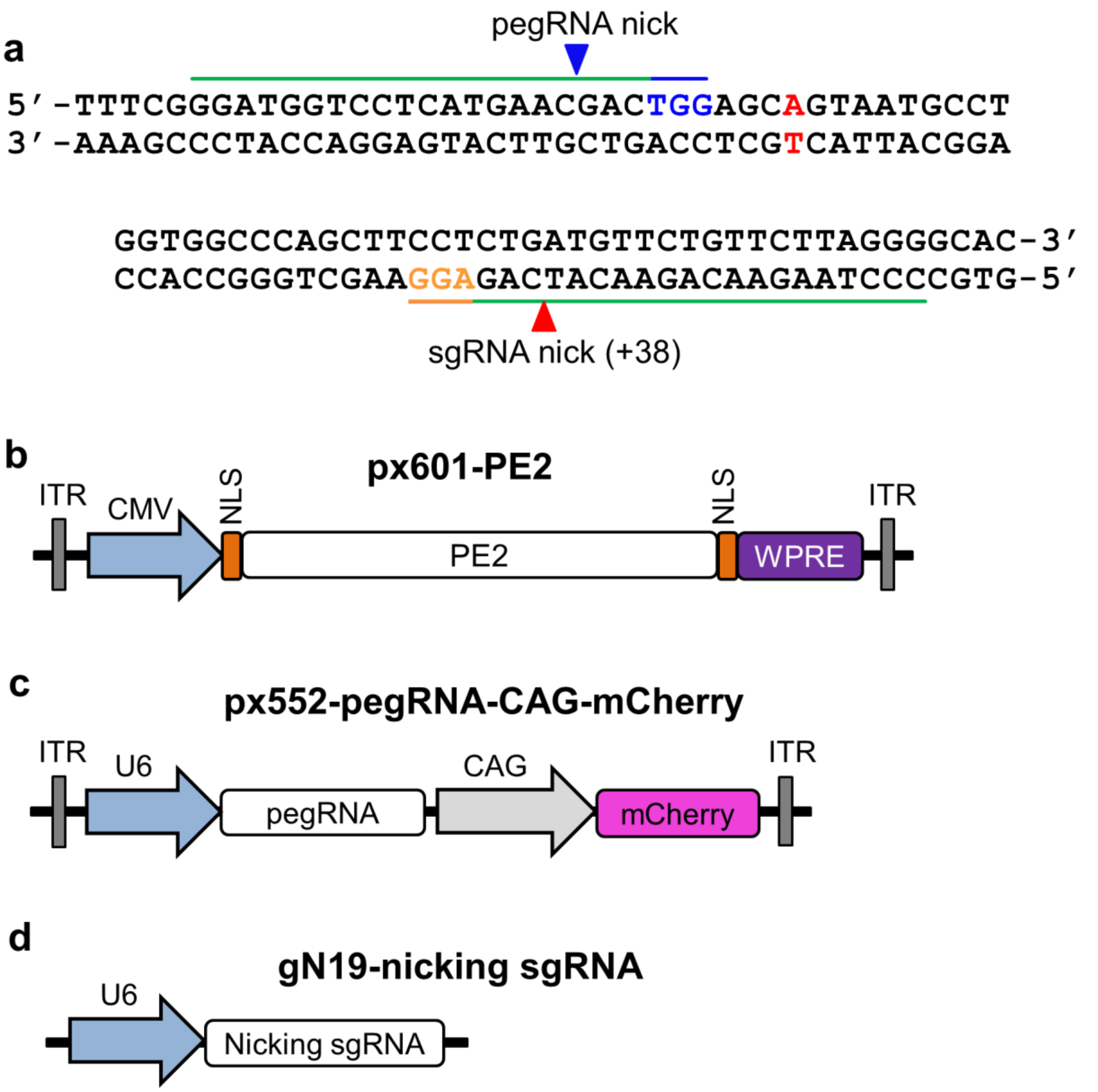


**Supplementary Figure 2. The target sequence in *Fah*$^{mut/mut}$ mice and maps of vectors encoding PE3 components used for treatment.** (**a**) The target and neighboring sequences. Green lines represent the pegRNA and sgRNA spacers. The PAMs of the pegRNA and sgRNA target sequences are highlighted in blue and orange, respectively. The disease-causing G>A point mutation is shown in red. (**b-d**) Vector maps of plasmids encoding PE2 (b), pegRNA (c), and sgRNA (d). ITR, inverted terminal repeat; CMV, cytomegalovirus promoter; NLS, nuclear localization sequence; WPRE, woodchuck hepatitis virus posttranscriptional regulatory element; CAG, CMV early enhancer/chicken β actin promoter; U6, human U6 promoter.

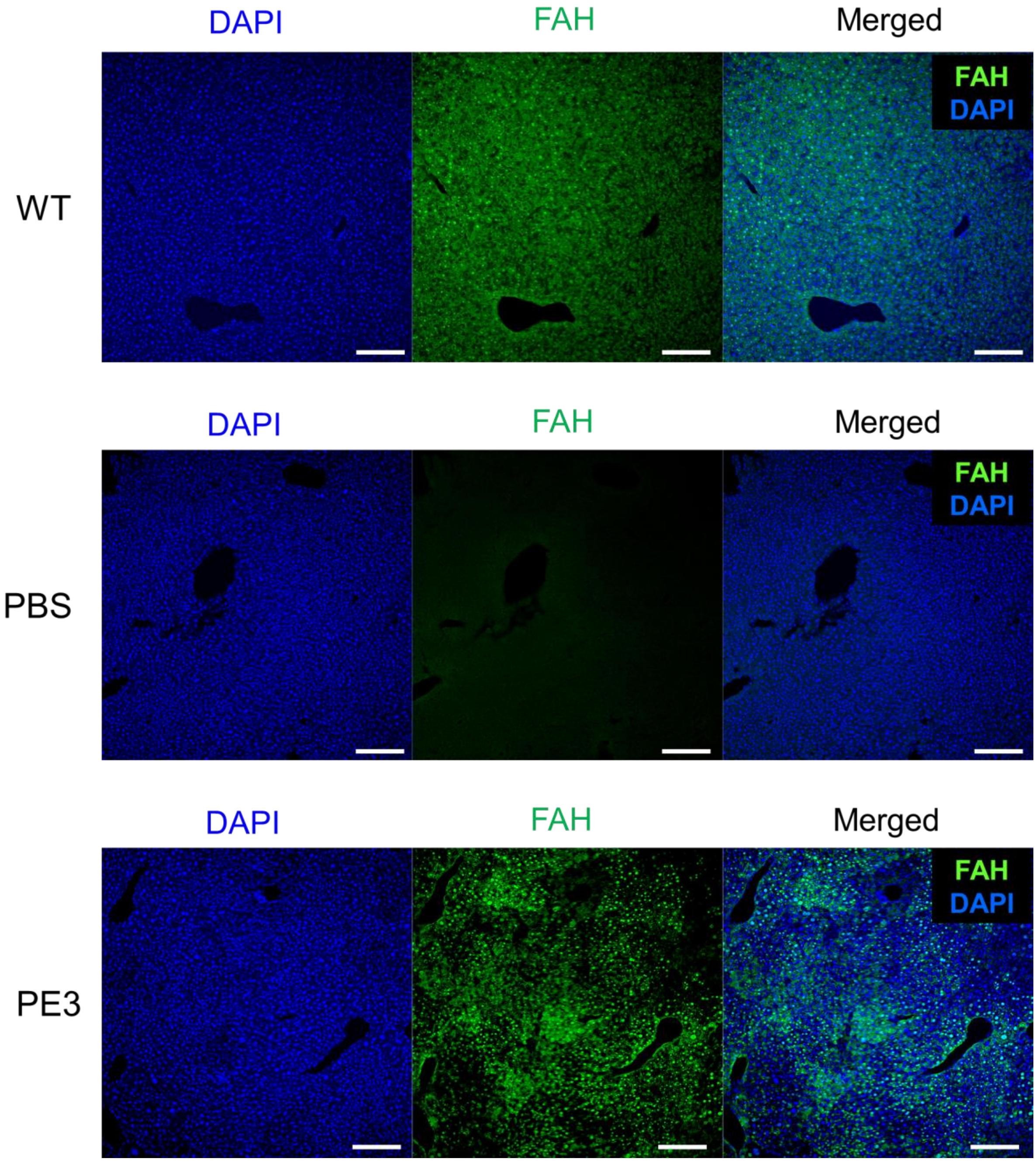


**Supplementary Figure 3.** Representative images from immunofluorescence experiments to detect the FAH protein in wild-type (WT), PBS-treated *Fah*$^{mut/mut}$ (PBS), and PE3-treated *Fah*$^{mut/mut}$ (PE3) mice. Merged images are shown on the right. Scale bar = 200 µm.

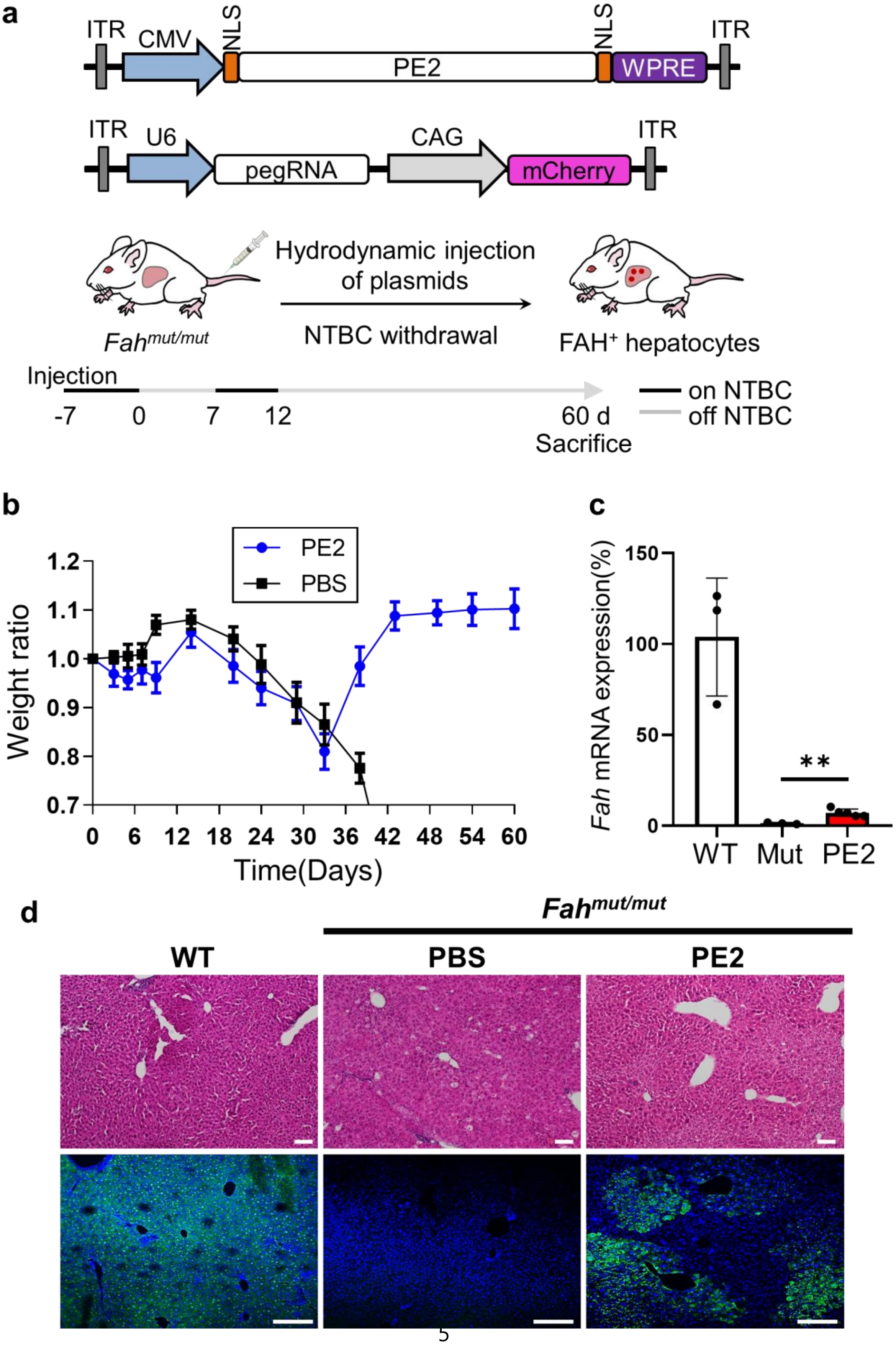

a
ITR
CMV
NLS
PE2
NLS
WPRE
ITR
ITR
U6
pegRNA
CAG
mCherry
ITR
Hydrodynamic injection of plasmids
NTBC withdrawal
Fah mut/mut
FAH+ hepatocytes
Injection
-7
0
7
12
60 d
Sacrifice
on NTBC
off NTBC
b
PE2
PBS
Weight ratio
Time(Days)
c
Fah mRNA expression(%)
**
WT
Mut
PE2
d
Fah mut/mut
WT
PBS
PE2

**Supplementary Figure 4. Prime editor 2 corrects the disease mutation and phenotype in *Fah*$^{mut/mut}$ mice.** (**a**) A schematic representation of the experiments. *Fah*$^{mut/mut}$ mice underwent injection of plasmids encoding prime editor 2 components (i.e., prime editor 2 and pegRNA) and were kept on water containing NTBC for 7 days. The day on which NTBC was withdrawn is defined as day 0. At 60 days, the PE2-treated mice were euthanized and analyzed. (**b**) Body weight of *Fah*$^{mut/mut}$ mice injected with PE2 or phosphate-buffered saline (PBS, control). Body weights were normalized to the pre-injection weight. The number of mice n = 5 for the PE2 group and n = 3 for the PBS group. Data are mean ± s.e.m. (**c**) The level of wild-type *Fah* mRNA in the liver measured by quantitative RT-PCR using primers that hybridize to exons 8 and 9. WT, wild-type mice; Mut, *Fah*$^{mut/mut}$ mice; PE2, *Fah*$^{mut/mut}$ mice injected with plasmids encoding PE2 components. The number of mice n = 3 (WT), 3 (Mut), and 5 (PE2). ***P* = 0.0032. (**d**) H&E staining (upper panels) and immunofluorescent staining for FAH protein (lower panels) in liver sections. Scale bars, upper panels, 100 μm; lower panels, 200 μm.

**Control Retina**

| | |
|---|---|
| GCCTTTCAGTTTCTGGGTGGGTGGTACTTCTAC | Unedited, 8,448 reads (99.90%) |
| GCCTTTCAGTTTCTGGATGGGTGGTACTTCTAC | 6 reads (0.07%) |
| GCCTTTCAGTTTCTGGTTGGGTGGTACTTCTAC | 2 reads (0.02%) |
| GCCTTTCAGTTTCTGGCTGGGTGGTACTTCTAC | 0 reads (0.00%) |

**AAV-PE3-injected Retina**

| | |
|---|---|
| GCCTTTCAGTTTCTGGGTGGGTGGTACTTCTAC | Unedited, 8,201 reads (97.92%) |
| GCCTTTCAGTTTCTGGATGGGTGGTACTTCTAC | 170 reads (2.03%) |
| GCCTTTCAGTTTCTGGTTGGGTGGTACTTCTAC | 4 reads (0.05%) |
| GCCTTTCAGTTTCTGGCTGGGTGGTACTTCTAC | 0 reads (0.00%) |

**Control RPE**

| | |
|---|---|
| GCCTTTCAGTTTCTGGGTGGGTGGTACTTCTAC | Unedited, 8,711 reads (99.82%) |
| GCCTTTCAGTTTCTGGATGGGTGGTACTTCTAC | 13 reads (0.15%) |
| GCCTTTCAGTTTCTGGTTGGGTGGTACTTCTAC | 2 reads (0.02%) |
| GCCTTTCAGTTTCTGGCTGGGTGGTACTTCTAC | 0 reads (0.00%) |

**AAV-PE3-injected RPE**

| | |
|---|---|
| GCCTTTCAGTTTCTGGGTGGGTGGTACTTCTAC | Unedited, 7,788 reads (98.50%) |
| GCCTTTCAGTTTCTGGATGGGTGGTACTTCTAC | 112 reads (1.42%) |
| GCCTTTCAGTTTCTGGTTGGGTGGTACTTCTAC | 4 reads (0.05%) |
| GCCTTTCAGTTTCTGGCTGGGTGGTACTTCTAC | 3 reads (0.04%) |

**Supplementary Figure 5.** Representative sequencing results from the retina and RPE of wild-type mice with AAV-PE3 injection and without injection (control). DNA sequences and the read count of sequences containing each mutation type are shown. Red type indicates the targeted position.

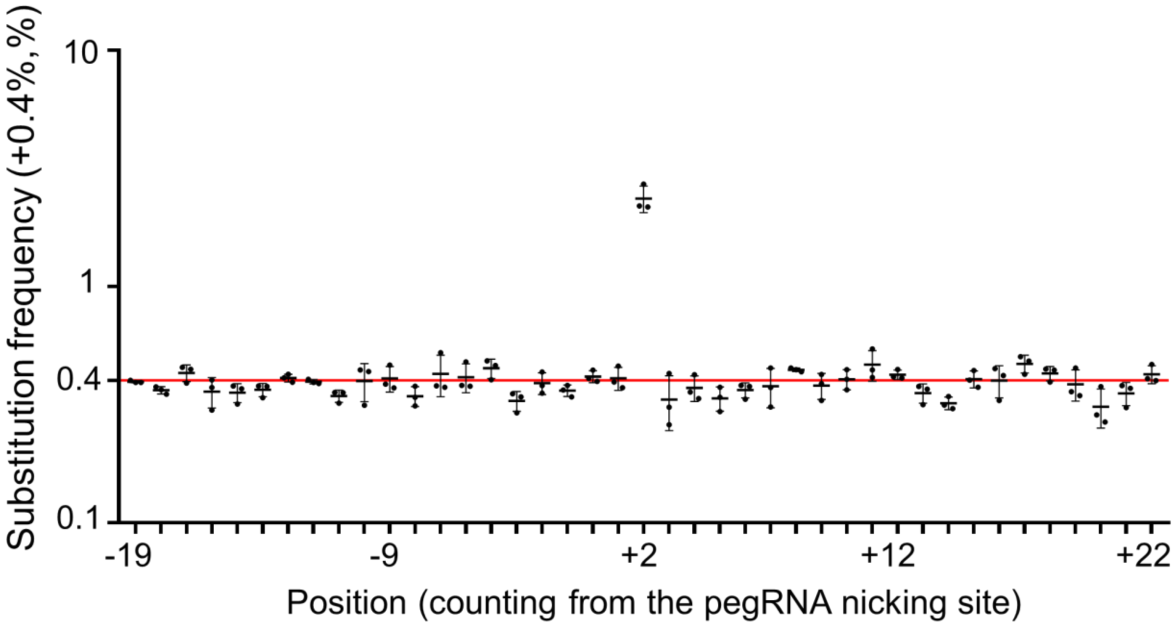


**Supplementary Figure 6.** Substitution frequencies at each position of the target sequence in PE3-treated retinas. The frequencies were normalized by subtracting the background substitution frequencies in controls that were not treated with PE3. Positions are numbered from the pegRNA nicking site. The targeted position is at +2. The red horizontal line represents the position where the normalized frequency = 0. Data are mean ± s.d. The number of mice n = 3.

**Supplementary Table 1**. Predicted activities of SpCas9 and SpCas9-NG at nine target sequences for the prime editing of the tyrosinemia-causing mutation (provided as a separate file).

**Supplementary Table 2**. Measured prime editing efficiencies for the tested pegRNAs using a paired library approach (provided as a separate file).

**Supplementary Table 3**. Predicted activities of the sgRNA candidates for the PE3-directed correction of the tyrosinemia-causing mutation (provided as a separate file).

**Supplementary Table 4**. Top-ranking potential off-target sites (provided as a separate file). The potential off-target sites were predicted by CRISPOR[17].

**Supplementary Table 5**. Predicted activities of the sgRNA candidates for the PE3-directed editing of *Atp7b* (provided as a separate file).

**Supplementary Table 6**. Comparison of efficiencies and precision of genome editing methods when genome editing tools were delivered using hydrodynamic injections in a mouse model of hereditary tyrosinemia (provided as a separate file).

**Supplementary Table 7**. Oligonucleotides used in our study (provided as a separate file).
